# Musicianship and active musical engagement predict facilitation of auditory-motor plasticity: evidence from auditory-motor paired associative stimulation

**DOI:** 10.1101/2025.10.21.683684

**Authors:** Shizuka Sata, Inhyeok Jeong, Naotsugu Kaneko, Sotaro Kondoh, Kimitaka Nakazawa, Shinya Fujii

## Abstract

**Background and Aim:** Auditory–motor paired associative stimulation (PAS) probes experience-dependent plasticity in auditory–motor circuits. We tested whether musical experience and musical sophistication modulate PAS-induced facilitation when instrument tones are paired with transcranial magnetic stimulation (TMS) over the left primary motor cortex.

**Method:** Sixteen musicians and thirteen non-musicians completed two sessions. Session 1 estimated an individualized inter-stimulus interval (ISI) by testing seven tone–TMS delays (25–300 ms). Session 2 applied PAS at the optimal ISI. Corticospinal excitability was assessed by motor-evoked potentials (MEPs) from right first dorsal interosseus at baseline and post-intervention. Group comparisons and a stepwise multiple regression with Goldsmiths Musical Sophistication Index (Gold-MSI) subscales evaluated predictors of the PAS effect.

**Results:** A stepwise multiple regression model selected “Group” and “Active Engagement” (Gold-MSI Factor 1) as independent predictors (*F* (2,26) = 4.59, *p* = 0.020, adjusted *R*^*2*^ = 0.204), without interaction: musicians exhibited smaller facilitation than non-musicians, and higher Active Engagement predicted greater facilitation across groups.

**Discussion:** The lack of facilitation in musicians contrasts with somatosensory PAS reports of enhanced plasticity in experts. A modality-specific ceiling effect may contribute, whereby long-term training optimizes auditory–motor transmission, reducing headroom for facilitation. The association with Active Engagement suggests motivational/reward mechanisms—potentially dopaminergic— gate responsiveness to auditory–motor PAS.

**Conclusion:** Under this protocol, auditory–motor PAS with instrument tones facilitated corticospinal excitability in non-musicians but not in musicians. Individual differences in Active Engagement predicted facilitation, indicating that both group membership and active musical engagement shape susceptibility to auditory–motor associative plasticity.

## 1 Introduction

The central nervous system exhibits remarkable flexibility, continuously adapting to experience and environmental demands through a process known as neuroplasticity. Musicians provide a valuable model for studying experience-dependent plasticity because their extensive, long-term training, which drives significant functional and structural adaptations in the brain [1, 2]. This training involves the precise, high-level integration of sensory information with fine motor commands, making musicians an informative population for probing mechanisms of adaptation.

One non-invasive method to investigate and induce neuroplasticity in humans is paired associative stimulation (PAS) [3]. The PAS protocol repeatedly pairs a sensory stimulus with a transcranial magnetic stimulation (TMS) pulse over the primary motor cortex (M1), thereby modulating synaptic efficacy through Hebbian-like mechanisms of long-term potentiation (LTP) [4] or depression (LTD) [5]. Previous studies using PAS combining somatosensory stimulation with M1-TMS have shown that musicians exhibit more pronounced bidirectional neuromodulation—both facilitatory (LTP-like) and inhibitory (LTD-like)—than non-musicians [6]. These findings can be interpreted as reflecting metaplasticity, whereby long-term musical training renders the musician’s brain more amenable to experience-dependent plasticity.

Building on the traditional somatosensory–motor focus of PAS, Sowman et al. developed an auditory–motor PAS protocol pairing speech sounds with M1-TMS [7], thereby enabling the investigation of auditory–motor plasticity. D’Ausilio et al. demonstrated that corticospinal excitability in pianists is facilitated in a manner dependent on the trained piece [8]. This finding suggests that musical experience and sophistication might shape auditory–motor plasticity. However, the extent to which individual differences in musical experience and musical sophistication predict the magnitude or direction of auditory–motor PAS effects remains unclear. Therefore, the present study aimed to investigate the influence of musical experience and sophistication on the efficacy of auditory–motor PAS. We applied an auditory–motor PAS protocol using instrument sounds in both musicians and non-musicians to test two questions: (i) whether PAS-induced changes in MEP amplitude differ between groups, and (ii) whether indices of musical sophistication predict inter-individual variability in PAS effects. Clarifying these relationships can refine models of auditory– motor learning and inform how prior experience shapes neuroplastic responses to non-invasive stimulation.

## 2 Materials and Methods

### Participants

Initially, 33 healthy volunteers were recruited. To account for potential dropouts, the initial number of participants was larger than that of previous studies comparing professional musicians and non-musicians [6]. Participants were assigned to one of two groups based on their musical background: a group of 16 highly trained musicians with over 10 years of piano experience (9 females) and a group of 17 non-musicians without formal education from a music school or conservatory (4 female) (see Table S1 for details of the participant characteristics described in this paragraph). Four participants from the non-musician group withdrew after the first experiment, leaving a final sample of 29 participants (16 musicians and 13 non-musicians) who completed all sessions. Handedness was assessed for all participants using the Edinburgh Handedness Inventory [9]. Musical sophistication was assessed using the Japanese version of the Goldsmiths Musical Sophistication Index (Gold-MSI) [10,11], which was developed to measure musical sophistication in both musician and non-musician.

The Gold-MSI consists of five subscales (Factor1: Active Engagement, Factor2: Perceptual Abilities, Factor3: Musical Training, Factor4: Singing Abilities, Factor5: Emotions). While the raw scores of subscales are reported in Table S2, they were z-scored for the analysis to standardize them, as they have different maximum scores.

All participants provided written informed consent. Eligibility was restricted to individuals with no contraindications for TMS. The study’s procedures were approved by the local ethics committee of the University of Tokyo (No. 754-7) and were conducted in accordance with the Declaration of Helsinki.

### Electromyography (EMG)

Surface EMG electrodes (Ag/AgCl; Vitrode F-150-S, Nihon Kohden, Japan) were placed on the right first dorsal interosseus (FDI). The interelectrode distance was 10 mm to prevent cross-talk between neighboring muscles. EMG signals were amplified 1000-fold and filtered with a band-pass filter of 5–1000 Hz (MEG-6108, Nihon Kohden, Japan). All EMG signals were digitized at 4000 Hz using an analog-to-digital converter (Power lab 4/26SP, PL2604, PLCI1, AD Instruments, Castle Hill, Australia) and stored on a local computer for offline analysis. EMG activity was monitored in real-time using data analysis software (LabChart 8, AD Instruments, Castle Hill, Australia).

### Auditory stimuli

A monaural piano tone corresponding to A3 was generated in Ableton Live (Version 11.3.42; Ableton AG, Berlin, Germany) using the factory instrument “Grand Piano”. A single MIDI note (A3; duration 3,000 ms) was rendered at a project sample rate of 44.1 kHz and exported to WAV format.

During the experiment, the audio was presented and hardware-timed using a Multi Trigger System MTS-0410 (Medical Try System, Japan), which issued a TTL trigger at acoustic onset each trial. Audio was delivered via insert earphones, held constantly across participants at 80dB SPL at the ear.

### Transcranial Magnetic Stimulation (TMS)

TMS was delivered to the left M1 using a figure-eight coil (70 mm external diameter) connected to a magnetic stimulator (Magstim 200, Magstim Co. Ltd., Whitland, UK), which produced monophasic pulses. We targeted the left M1, for which a structural left-hemisphere advantage has been reported in pianists [12]. For the paired-stimulus conditions, the stimulator was triggered externally by TTL pulses from the Multi Trigger System to ensure precise temporal synchronization with the auditory stimuli. During the experiment, participants were seated comfortably in a reclining chair with their arms supported by armrests. The coil was held tangentially to the scalp with the handle pointing posteriorly and laterally at approximately 45° to the sagittal midline, an orientation known to induce a posterior-to-anterior current flow.

The optimal coil position, “hotspot”, was identified as the left M1 site that consistently elicited the largest MEPs from the contralateral FDI muscle. This location was registered with a TMS tracking system (BrainSight, Rogue Research, Montreal, QC, Canada) to ensure precise and consistent coil placement throughout the experiment. TMS pulses were delivered with a randomized inter-trial interval of 5,500 ± 1,000 ms. During all TMS procedures, EMG activity of the target muscle was monitored in real-time to ensure it remained at rest prior to stimulation. Trials showing pre-stimulus EMG activity exceeding 10 µV within the 100 ms preceding the pulse were excluded from analysis; when exclusions occurred, replacement trials were acquired immediately to maintain the planned number of MEPs.

To account for inter-individual variability in corticospinal excitability, the TMS intensity was normalized for each participant by standardizing the resulting MEP amplitude. At the beginning of each experiment, the stimulator output was individually adjusted to determine the intensity required to elicit MEPs with a peak-to-peak amplitude of approximately 1 mV (Stimulus Intensity 1mV: SI1mV) [6]. This procedure ensured that the physiological baseline was comparable across all participants before each experiment. In addition, we asked participants to abstain from caffeine consumption on the day of the experiment to avoid any potential interference with baseline corticospinal excitability [13].

### Auditory-Motor Paired Associative Stimulation (Auditory-Motor PAS)

The experiment consisted of two sessions separated by an interval of at least 7 but no more than 31 days. This separation was necessary to analyze data from the first session and determine the optimal, individualized inter-stimulus interval (ISI) for the subsequent PAS intervention. We chose an individualized approach because our preliminary experiments revealed no single ISI that consistently facilitated MEPs across all individuals. The seven ISIs tested (25, 50, 100, 150, 200, 250, and 300 ms) were selected based on those used by a previous study of auditory-motor PAS [7]. First, a baseline measurement of corticospinal excitability was established by recording 16 MEPs from single-pulse TMS. During this baseline block, participants wore insert earphones with no sound delivered to create a silent condition. Following the baseline recording, MEPs were elicited by pairing an auditory stimulus with a TMS pulse at these seven ISIs. Sixteen paired stimuli were delivered for each ISI condition. To counteract order effects, the presentation sequence of these seven conditions was randomized, with the same sequence used for all participants to ensure comparability between groups. For each participant, the optimal ISI was defined as the interval that produced the largest mean MEP amplitude.

In the second session, an auditory-motor PAS protocol was applied using the individualized ISI. The intervention consisted of 200 pairs of auditory and TMS stimuli. However, for two non-musician participants, PAS was performed using the ISI that had elicited the second-highest mean MEP due to a calculation error in identifying the optimal interval. Corticospinal excitability was assessed by recording 15 MEPs in a silent condition at four time points: before the intervention (Baseline), and at 0, 15, and 30 minutes post-intervention. To control potential circadian influences on neural excitability [14], the time of day for this PAS session (morning or afternoon) was balanced between the two groups.

### Data processing and statistical analysis

Group comparability for age, sex, and TMS stimulus intensity were assessed using an unpaired t-test (Welch’s correction when appropriate) and Fisher’s exact test (two-sided), respectively. The TMS stimulus intensity (SI1mV) was expressed as % of maximum stimulator output (%MSO).

MEPs for each ISI were analyzed using a mixed repeated-measures ANOVA with ISI (7 levels: 25, 50, 100, 150, 200, 250, and 300 ms) as a within-subject factor and Group (2 levels: Musician and Non-musician) as a between-subject factor. Sphericity was assessed with Mauchly’s test. *p*-values set to 0.05 were corrected using the Bonferroni method.

To quantify the overall effects of the intervention effect as a single representative value, we first calculated the mean MEP amplitude from the three post-intervention time points (0, 15, and 30 min) to yield a single “post-PAS” value for each participant. This post-PAS value was then normalized to the individual’s baseline amplitude and expressed as a percentage (e.g., baseline = 100%). To assess the within-group effect of PAS, a one-sample t-test was conducted for each group to determine if the normalized post-PAS MEP amplitude was significantly different from 100 (%). Because we specifically hypothesized a facilitatory effect (an increase from 100%) [7], these t-tests were one-tailed.

To investigate whether musical experience (group) and sophistication could predict PAS-induced plasticity, a multiple linear regression analysis with stepwise variable selection (forward and backward) based on minimizing the Bayesian Information Criterion (BIC) was performed. The dependent variable was the post-PAS MEP percentage change rate. The initial set of predictor variables (candidates) included Group (musician vs. non-musician) and the z-scored five subscales of the Gold-MSI. The categorical variable ‘Group’ was sum-coded (Musicians = +1, Non-Musicians = −1); the group coefficient therefore represents half the Musicians–Non-Musicians difference. The model examined the main effects of the candidate predictors and their interaction effects. Multicollinearity among the predictor variables was assessed by calculating the Variance Inflation Factor (VIF).

Data processing and statistical analyses were performed using MATLAB (R2025b; MathWorks, MA, USA), SPSS Statistics (Version 30.0.0; IBM Corp., Armonk, NY, USA), and R (Version 4.3.2; R Foundation for Statistical Computing, Vienna, Austria).

## 3 Results

### Group characteristics

There were no significant differences between the two groups in age, sex, handedness, or TMS stimulus intensity (see Table S2). For the ISI, no conditions showed significant differences either within-group or between-groups (see Table S3).

### Auditory-Motor Paired Associative Stimulation (Auditory-Motor PAS)

Post-PAS MEPs (expressed as % of baseline) of Within-group one-sample tests against 100% showed a significant deviation in non-musicians but not in musicians (non-musicians: *t* (12) = 2.14, *p* = 0.027, *g* = 0.56; musicians: *t* (15) = 0.63, *p* = 0.538, *g* = 0.15) (Figure1.A).

**Figure 1.**
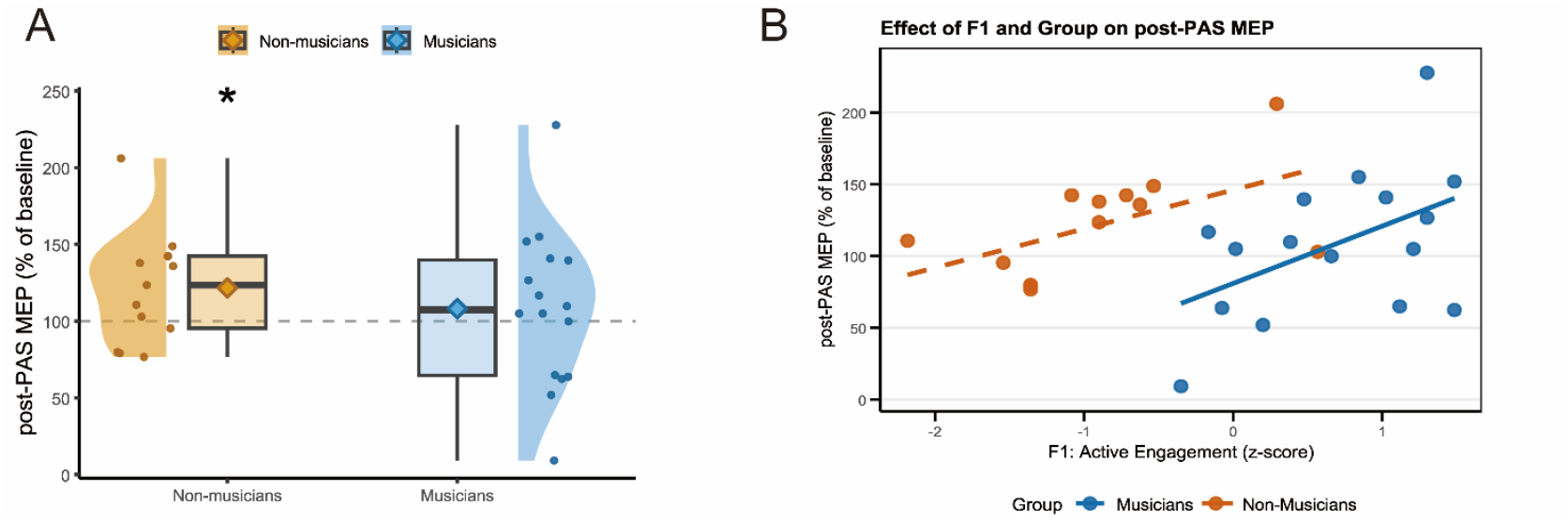
Relationship between Auditory-Motor PAS effect and Musical experience. (A) Violin–box plots show post-PAS motor evoked potential (MEP) amplitudes expressed as a percentage of baseline (100%) in non-musicians (orange) and musicians (blue). Each small circle represents an individual participant. The central box shows the interquartile range (IQR; 25th–75th percentiles), and the horizontal line inside the box indicates the median. The surrounding violin shapes represent the kernel density of the data distribution. Diamonds indicate the group mean values. Non-musicians exhibited a significant increase in post-PAS MEP relative to baseline (\**p* < 0.05), whereas musicians showed no significant change. (B) Scatter plot illustrating the association between Active Engagement (Gold-MSI F1, z-scored) and post-PAS MEP (% of baseline). Points are individual participants; separate regression lines are shown for musicians (blue) and non-musicians (orange). The stepwise multiple regression model selected Group and F1 as predictors, and the Group × F1 interaction was not significant.

To identify predictors of Auditory-Motor PAS effects, we conducted a multiple linear regression analysis with the post-PAS MEP amplitude as the dependent variable. A stepwise variable selection procedure using the BIC resulted in a final model that included Factor1 (Active Engagement) and group as predictors. The final model was statistically significant (*F* (2, 26) = 4.59, *SE* = 11.45, *p* = 0.020, Adjusted *R*^*2*^ = 0.204). Key assumption checks did not provide evidence against normality of residuals (Shapiro-Wilk test, *W* = 0.96, *p* = 0.42) and VIF was 2.45, suggesting that multicollinearity was unlikely to bias parameter estimates severely. The analysis revealed that both F1 (*Estimate* = 33.16, *SE* = 11.78, *p* = 0.008) and group (*Estimate* = −32.98, *p* = 0.009) were significant predictors (see Figure1.B). This indicates that higher Active Engagement was associated with greater facilitation of MEP amplitude after PAS. In contrast, the musician group exhibited lower post-PAS MEP amplitudes than the non-musician group.

## 4 Discussion

This study aimed to test whether musical experience and sophistication modulate the facilitatory effect of auditory–motor PAS using instrument tones. Under the present protocol, t-tests showed a significant increase in MEP amplitude following PAS in non-musicians, but not in musicians. In addition, an exploratory multiple regression indicated that both Group and the “Active Engagement” (Factor1) subscale accounted for additional variance: musicians showed a smaller increase in MEP amplitude than non-musicians, whereas higher Active Engagement predicted greater facilitation, independent of Group.

We found that auditory-motor PAS did not induce significant MEP facilitation in musicians, whereas it was significantly induced in non-musicians. These results differed from previous research of somatosensory PAS, which showed a more pronounced effect in musicians than in non-musicians [6]. The groups did not significantly differ in age, sex, handedness, or TMS stimulus intensity (i.e., baseline MEP), suggesting that these factors cannot account for the observed group difference in MEP facilitation. This discrepancy may be due to the difference in sensory modality. Specifically, while long-term musical training may enhance the plasticity of somatosensory–motor circuits, it may also lead to an optimization—or saturation—of auditory–motor pathways [1]. The robust audio-motor neural pathways established through extensive musical training may therefore have been less susceptible to the influence of PAS. In such cases, the capacity for further plastic changes is limited, a phenomenon known as a ceiling effect. This idea is supported by a study using transcranial direct current stimulation, in which improvements in fine motor control during metronome-synchronized keystrokes were observed in non-musicians but not in expert pianists [15]. The absence of PAS-induced facilitation in the auditory–motor system of musicians may thus reflect a lack of available plasticity rather than an absence of functional connectivity. By contrast, even in highly trained individuals, the somatosensory–motor system may retain greater capacity for facilitation, suggesting that the extent of plasticity induced by PAS can depend on both the sensory modality and the degree of prior use-dependent optimization.

A stepwise multiple regression analysis revealed that both Group and Active Engagement were significant predictors of the auditory-motor PAS effect. The finding that the musician group was less modulated by the PAS effect is consistent with the above description. Active Engagement is a factor that reflects an attitude of active involvement with music through the investment of resources such as time and money, including concert attendance and a high sensitivity to musical exploration [10]. The link between Active Engagement and PAS effectiveness may point to the role of the brain’s reward system, which is underpinned by dopaminergic neural circuits. It has been shown that individuals with higher subjective pleasure ratings and greater reward-related brain activity are more willing to purchase music [16]. This reward system is enhanced by the administration of levodopa, a dopamine precursor, which increases both the hedonic experience of and motivation for music [17]. Critically, levodopa has also been shown to modulate the effects of PAS [18, 19]. Therefore, our results suggest that individual differences in the sensitivity of dopaminergic circuits for musical motivation and reward may be involved. The activation of this reward system through Active Engagement may thus increase the potential for plasticity.

## 5 Acknowledgements

We are grateful to all participants. We thank Mr. K. Ueda for creating the auditory stimuli, and Dr. I. Ayase and Ms. Y. Sakakibara for their advice on the statistical models.

---

**Table S1.**
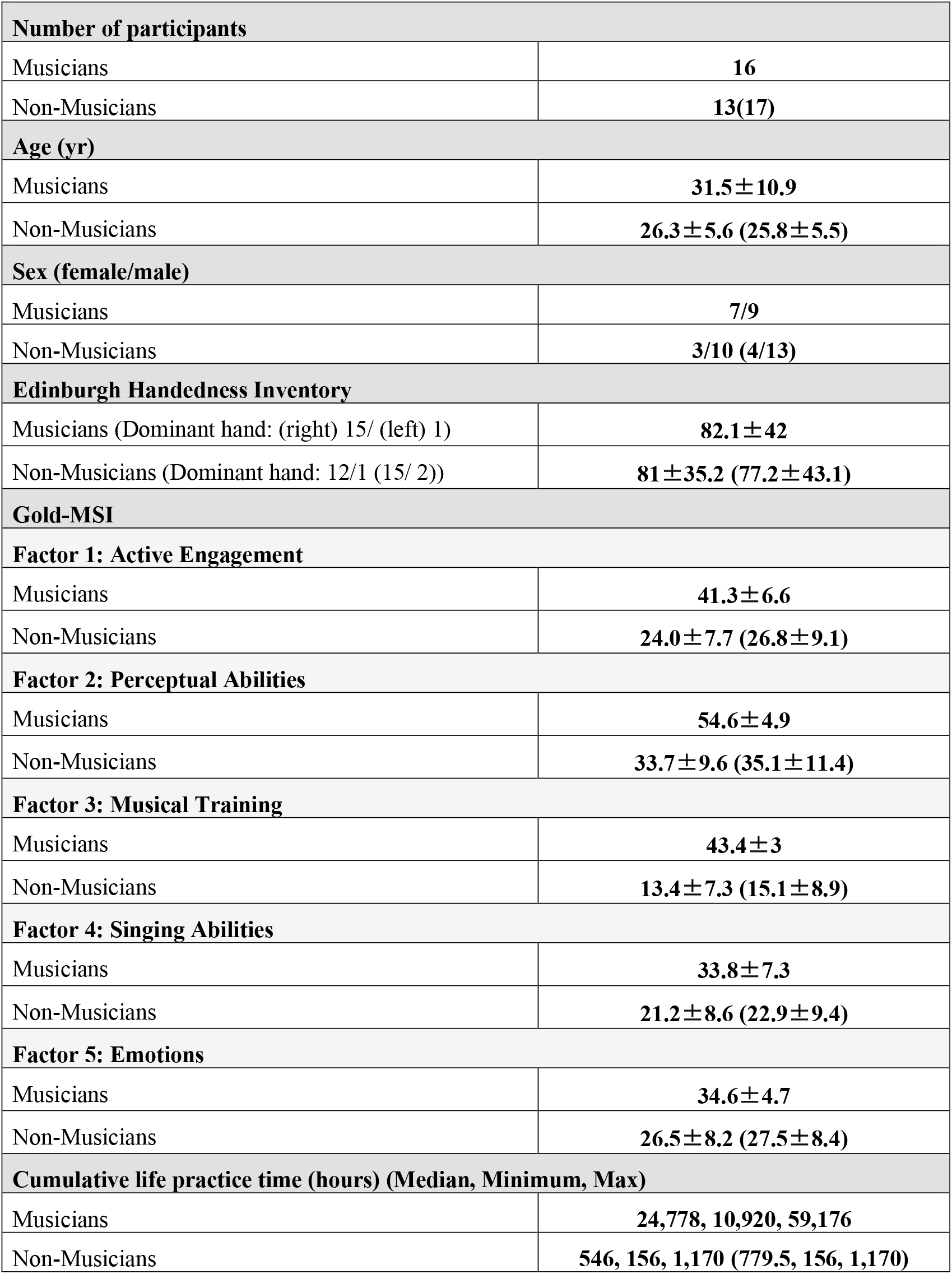
Participant characteristics *Note:* Non-Musicians (ISI+PAS (ISI only)), ± = SD.

| **Number of participants** |  |
| --- | --- |
| Musicians | 16 |
| Non-Musicians | 13(17) |
| **Age (yr)** |  |
| Musicians | 31.5±10.9 |
| Non-Musicians | 26.3±5.6 (25.8±5.5) |
| **Sex (female/male)** |  |
| Musicians | 7/9 |
| Non-Musicians | 3/10 (4/13) |
| **Edinburgh Handedness Inventory** |  |
| Musicians (Dominant hand: (right) 15/ (left) 1) | 82.1±42 |
| Non-Musicians (Dominant hand: 12/1 (15/ 2)) | 81±35.2 (77.2±43.1) |
| **Gold-MSI** |  |
| **Factor 1: Active Engagement** |  |
| Musicians | 41.3±6.6 |
| Non-Musicians | 24.0±7.7 (26.8±9.1) |
| **Factor 2: Perceptual Abilities** |  |
| Musicians | 54.6±4.9 |
| Non-Musicians | 33.7±9.6 (35.1±11.4) |
| **Factor 3: Musical Training** |  |
| Musicians | 43.4±3 |
| Non-Musicians | 13.4±7.3 (15.1±8.9) |
| **Factor 4: Singing Abilities** |  |
| Musicians | 33.8±7.3 |
| Non-Musicians | 21.2±8.6 (22.9±9.4) |
| **Factor 5: Emotions** |  |
| Musicians | 34.6±4.7 |
| Non-Musicians | 26.5±8.2 (27.5±8.4) |
| **Cumulative life practice time (hours) (Median, Minimum, Max)** |  |
| Musicians | 24,778, 10,920, 59,176 |
| Non-Musicians | 546, 156, 1,170 (779.5, 156, 1,170) |

**Table S2.** Participant demographics and stimulation intensity.

| Variable |  | Non-musicians | Musicians | Test statistic | p-value | Effect size (95% CI) |
| --- | --- | --- | --- | --- | --- | --- |
| Age (years) | ISI | 25.8 ± 5.7 | 31.5 ± 11.2 | Welch's t (21.84) = -1.83 | 0.081 | Hedges' g = -0.63 [-1.31, 0.06] |
|  | PAS | 26.3 ± 5.8 |  | t (27) = -1.51 (Levene p = 0.087) | 0.144 | g = -0.55 [-1.27, 0.19] |
| Sex (F/M) | ISI | 4 / 13 | 9 / 7 | Fisher's exact = 0.080; $\chi^2$ (1) = 3.70 | 0.080 | - |
| | PAS | 3 / 10 | | Fisher's exact = 0.130; $\chi^2$ (1) = 3.25 | 0.130 | |
| SI1mV (%MSO, ISI) |  | 72.9 ± 13.2 (n=17) | 71.1 ± 12.6 (n=16) | t (31) = 0.40 | 0.689 | g = 0.14 [-0.53, 0.80] |
| SI1mV (%MSO, PAS) |  | 72.0 ± 11.2 (n=13) | 68.6 ± 10.6 (n=16) | t (28) = 0.85 | 0.404 | g = 0.30 [-0.40, 1.00] |

*Note.* SI1mV = stimulus intensity required to evoke ~1 mV MEP; %MSO = percentage of maximum stimulator output.

**Table S3.**
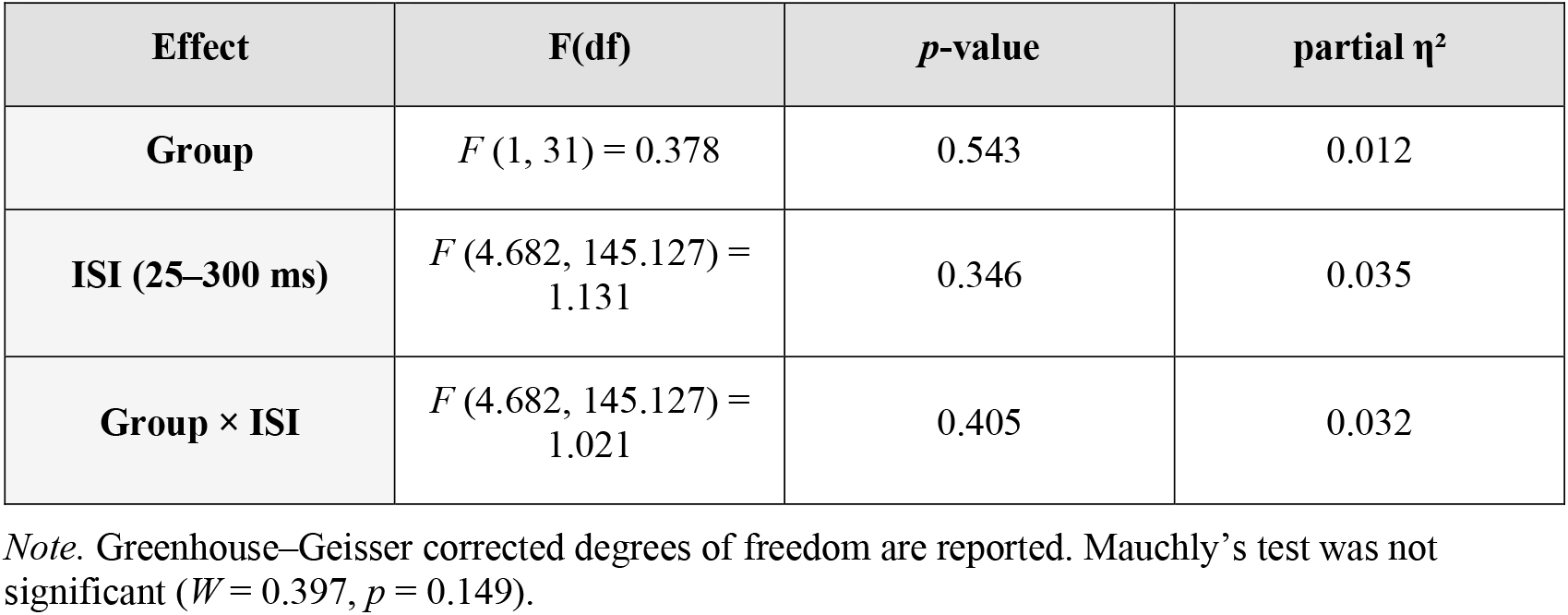
Mixed-design ANOVA results for ISI experiment.

| Effect | F(df) | *p*-value | partial $\eta^2$ |
| --- | --- | --- | --- |
| Group | $F(1, 31) = 0.378$ | 0.543 | 0.012 |
| ISI (25–300 ms) | $F(4.682, 145.127) = 1.131$ | 0.346 | 0.035 |
| Group × ISI | $F(4.682, 145.127) = 1.021$ | 0.405 | 0.032 |

*Note.* Greenhouse–Geisser corrected degrees of freedom are reported. Mauchly’s test was not significant ( $W = 0.397, p = 0.149$ ).

